# Interleukin 11 expression causes murine inflammatory bowel disease

**DOI:** 10.1101/756098

**Authors:** Wei-Wen Lim, Benjamin Ng, Anissa Widjaja, Chen Xie, Liping Su, Nicole Ko, Sze-Yun Lim, Xiu-Yi Kwek, Stella Lim, Stuart A Cook, Sebastian Schafer

## Abstract

Interleukin 11 (IL11) is a profibrotic cytokine, secreted by myofibroblasts and damaged epithelial cells. Smooth muscle cells (SMCs) also secrete IL11 under pathological conditions and express the IL11 receptor. Here we examined the effects of SMC-specific, conditional expression of murine IL11 in a transgenic mouse (*Il11*^SMC^). Within days of transgene activation, *Il11*^SMC^ mice developed loose stools and progressive bleeding and rectal prolapse, which was associated with a 65% mortality by two weeks. The bowel of *Il11*^SMC^ mice was inflamed, fibrotic and had a thickened wall, which was accompanied by activation of ERK and STAT3. In other organs, including heart, lung, liver, kidney and skin there was a phenotypic spectrum of fibro-inflammation, together with consistent ERK activation. To investigate further the importance of stromal-derived IL11 in the inflammatory bowel phenotype we used a second model with fibroblast-specific expression of IL11, the *Il11*^Fib^ mouse. This additional model largely phenocopied the *Il11*^SMC^ bowel phenotype. These data show that IL11 secretion from the stromal niche is sufficient to drive inflammatory bowel disease in mice. Given that IL11 expression in colonic stromal cells predicts anti-TNF therapy failure in patients with ulcerative colitis or Crohn’s disease, we suggest IL11 as a therapeutic target for inflammatory bowel disease.

## Main text

Non-striated smooth muscle cells (SMCs) line the walls of hollow organs and the vasculature. In adults, SMCs are not terminally differentiated and their cellular phenotype remains plastic. A variety of extracellular cues such as humoral factors, mechanical or oxidative stress and cell-cell interactions can induce a spectrum of cellular states ranging from contractile SMCs to highly synthetic and proliferative SMCs [1]. Synthetic SMCs are associated with a wide variety of vascular pathologies such as atherosclerosis or hypertension [1] and other disorders such as asthma [2] and inflammatory bowel disease (IBD) [3]. Many fibro-inflammatory diseases have a component, or are defined by, SMC dysfunction. This is exemplified by systemic sclerosis, which presents with global organ fibrosis and specific vascular abnormalities [4] and is characterized by elevated transforming growth factor beta (TGFB) 2 and interleukin 11 (IL11) expression in dermal stromal cells [5,6]. This co-occurrence of fibrosis and SMC dysfunction may in part be explained by molecular similarities of the fibrogenic fibroblast-to-myofibroblast conversion and the SMC contractile-to-synthetic phenotype switch. Both these cellular transitions are characterized by extracellular matrix (ECM) production, cell proliferation, invasion and migration. They can also be triggered by the same extracellular cues including TGFB family members [1,7].

We recently identified IL11 as a critical driver of fibroblast activation in the cardiovascular system, liver and lung downstream of a variety of pro-fibrotic factors including TGFB1 [8–10]. In a study from 1999, IL11 was also found to be secreted by vascular SMCs (VSMCs) in response to pathogenic stimuli, including interleukin 1 alpha (IL1A), TGFB and tumor necrosis factor (TNF) [11]. Although IL11 is upregulated in systemic sclerosis [6], TNF-resistant ulcerative colitis [12,13] and asthma [14] and despite SMCs being a source of IL11 [11], the effect of IL11 function in SMC biology has not been studied. To address this gap in our knowledge, we generated an inducible *Il11* transgenic mouse to overexpress mouse *Il11* in myosin heavy chain 11 (*Myh11*)-positive smooth muscle cells (*Il11*^SMC^). Here we characterized key organs that may be affected by SMC pathobiology in *Il11*^SMC^ mice to better understand the role of SMC-derived IL11.

## Results

### Expression of Il11 in smooth muscle cells results in ill health and early mortality

We generated the *Il11*^SMC^ mouse model that overexpresses IL11 specifically in *Myh11*^*+ve*^ smooth muscle cells: Conditional transgenic mice with mouse *Il11* inserted into the Rosa26 locus (*Rosa26-Il11-*Tg) [8] *were crossed with smooth muscle-specific Myh11-cre/ERT2* mice [15] (**Fig. 1a, b**). We then injected tamoxifen (tam) three times at day 0, 3 and 5 into 6-week old *Il11*^SMC^ mice to induce recombination in *Myh11*^*+ve*^ cells and monitored survival and body weight for 14 days. Following tam-induced *Il11* expression in SMCs, mice started dying from day three onwards, with only 37% of *Il11*^SMC^ mice surviving to day 14. This was significantly different from the survival of either vehicle-treated *Il11*^SMC^ animals or tam-treated *Cre*^SMC^ control mice, which were unaffected and both had 100% survival (both P < 0.001; **Fig 1d** and **Supplementary Fig 1b**). Starting from day four onwards, tam-treated *Il11*^SMC^ mice progressively lost weight as compared to vehicle-treatment and tam-treated *Cre*^SMC^ controls (both P < 0.001; **Fig 1e** and **Supplementary Fig 1d)**. Following two weeks of tam-induced *Il11* expression, *Il11*^SMC^ mice were significantly smaller in body weight and length as compared to tam-treated *Cre*^SMC^ controls (both P < 0.001; **Supplementary Fig 1f, g**) and vehicle-treated *Il11*^SMC^ mice (P = 0.002 and P < 0.001 respectively; **Fig 1f, g**). In contrast, the indexed weight of the heart, lung and kidney in tam-treated *Il11*^SMC^ animals was significantly elevated (P_Heart_ < 0.001; P_Lung_ < 0.001; P_Kidney_ = 0.006) when compared to vehicle treated mice (**Fig 1h**). We did not observe differences in liver weight or colon length in veh or tam treated *Il11*^SMC^ animals (data not shown).

**Figure 1.**
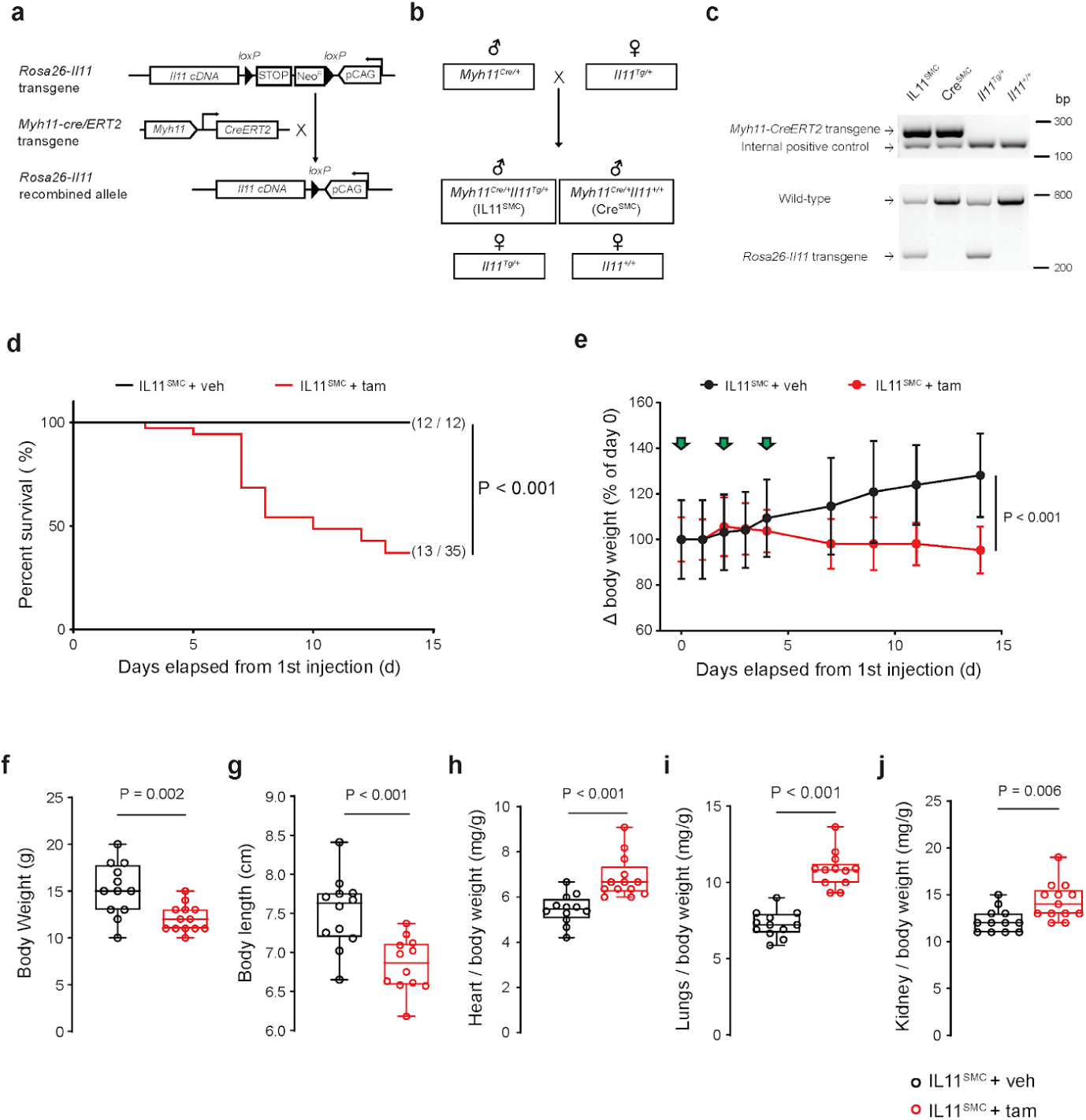
Expression of *Il11* in smooth muscle cells is associated with body weight loss, elevated organ weights and spontaneous death. **a** Schematic diagram of the targeted expression of *Il11* in *Myh11*^+ve^ SMC. In *Rosa26-Il11* mice, a floxed cassette containing both the neomycin (neo) resistance and stop elements is positioned before the murine *Il11* transgene cassette, which undergoes tamoxifen (tam) initiated *Cre*-mediated recombination when crossed to the *Myh11-Cre/ERT2* mouse. **b** Breeding scheme to generate *Myh11*^*Cre/+*^*Rosa26*^*Il11/+*^ (*Il11*^SMC^) and *Myh11*^*Cre/+*^*Rosa26*^*+/+*^ (*Cre*^SMC^) offspring mice. Note that the *Myh11-Cre* gene is expressed on the Y chromosome and therefore only male offspring carry the transgene. **c** Genotyping of tail biopsy DNA. A 287 bp band indicates the presence of the *Cre* transgene whereas the 180 bp band determines the presence of the internal positive control (top gel). Polymerase chain reaction with the *Rosa26-Il11* primer set detects a 270 bp band indicative of the *Rosa26-Il11* transgene whereas the 727 bp band indicates the presence of the wild-type transgene (bottom gel). **d** Survival curve of *Il11*^SMC^ mice treated with tam (*n* = 35) and corn oil vehicle (veh; *n* = 12) mice following tamoxifen initiation at day 0 and followed until day 14. Survival curves were compared using the log-rank Mantel-Cox test. **e** Body weight changes (expressed as percentage of day 0 body weight) in *Il11*^SMC^ mice treated with tam or veh (*n* = 8 per group). Green arrows denote individual injections. Statistical analyses by two-way ANOVA with Sidak multiple comparisons; data expressed as mean ± standard deviation. **f-g** Collated body weights (left) and body lengths (right) of *Il11*^SMC^ mice treated with tam or veh measured at d14 post initial tamoxifen dose (*n* = 12-13 per group). **h-j** Organ weights of the heart, lung and kidney normalized to body weight in *Il11*^SMC^ mice treated with tam or veh (*n* = 12-13 per group). All comparisons were conducted in mice 14 days post-veh and tam treatment. Statistical analyses by two-tailed unpaired t-test; data expressed as median ± IQR, whiskers representing the minimum and maximum values.

### Il11 expression causes severe inflammatory bowel disease associated with fibrosis

The most obvious and striking feature of *Il11*^SMC^ mice treated with tam was progressive rectal prolapse and pale loose stool formation from as early as day three after gene induction (**Fig 2a** and **Supplementary 1c**). Gross anatomical inspection of the gastrointestinal tract revealed injection and swelling of the small and large intestines of tam-treated *Il11*^SMC^ mice when compared to veh-treated controls (**Fig 2b**). Intestinal inflammation was specifically indicated by an increase in fecal calprotectin, a biomarker used to monitor disease activity in human colitis, of tam-treated *Il11*^SMC^ mice when compared to veh treatment (P < 0.001; **Fig. 2b, c**). Masson’s trichrome staining of the colon indicated a very large increase in collagen deposition (P < 0.001; **Fig 2d, e**). Histology also showed a significant increase in the thickness of the smooth muscle-dominant muscularis propria (P = 0.040; **Fig 2f**). Quantitative hydroxyproline assessments revealed an increase in colonic collagen content in *Il11*^SMC^ mice after tam treatment (P < 0.001; **Fig 2g**), confirming the histological data.

**Fig 2.**
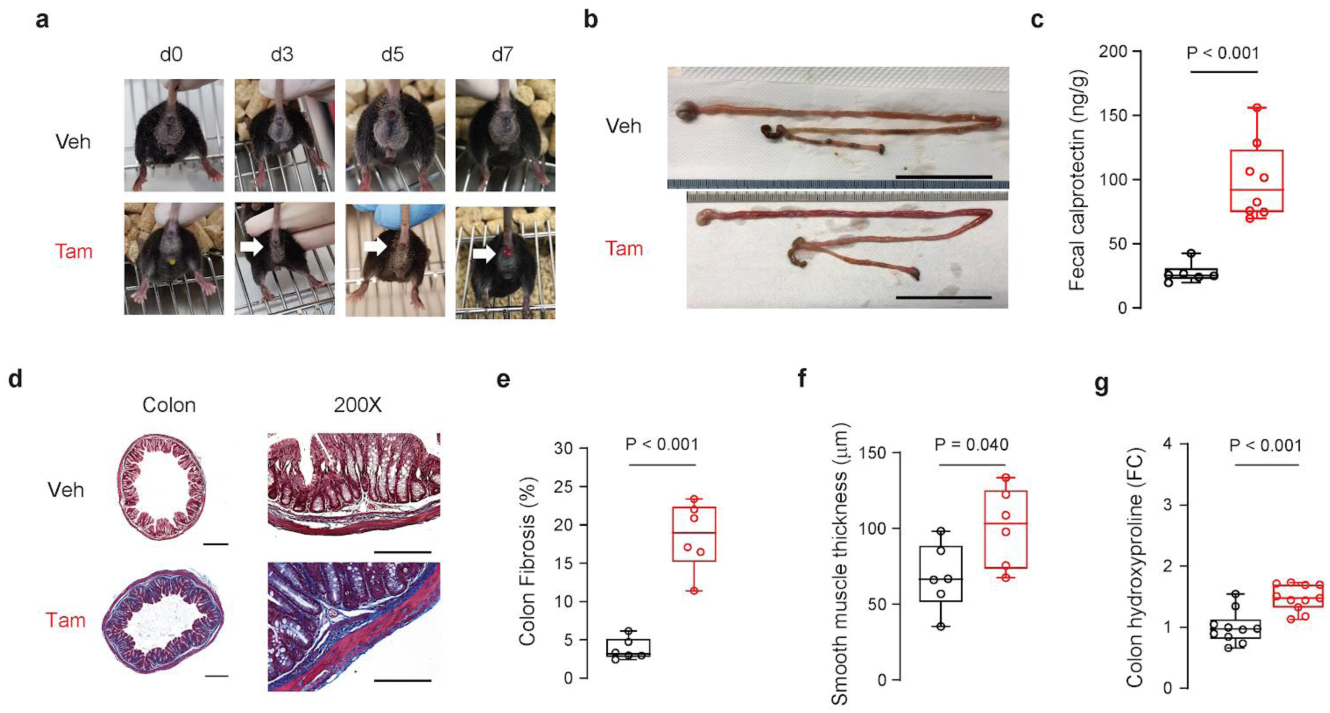
*Il11* expression results in fibro-inflammatory disease of the colon. **A** Representative images of the *Il11*^SMC^ mice before (d0) and up to 7 days (d7) treatment with either corn oil vehicle (veh) or tamoxifen (tam). Presence of rectal prolapse are indicated with white arrows. Images represented the same animal across time points not taken to the same scale. **b** Excised gastrointestinal tract of representative *Il11*^SMC^ mice at day 14 post-treatment with veh or tam. Scale bar represents 5 cm. **c** Fecal calprotectin in representative *Il11*^SMC^ mice treated with veh or tam assessed by ELISA (*n* = 6-8 per group). **d** Representative cross-section of the colon stained with Masson’s trichrome (left) and at 200X magnification (right). Scale bar of cross section represents 500 µm and 200X magnification represents 200 µm. **e** Colon fibrosis determined as a percentage of collagen positive area (blue) from histological images taken at 200X magnification (*n* = 6 per group). **f** Tunica muscularis (smooth muscle) thickness of the colon (*n* = 6 per group). **g** Total collagen content assessed by hydroxyproline assay and expressed as fold change (FC) of veh-treated *Il11*^SMC^ mice (*n* = 10-11 per group). All comparisons were conducted in organs harvested from mice 14 days post-veh and tam treatment. Statistical analyses by two-tailed unpaired t-test; data expressed as median ± IQR, whiskers representing the minimum and maximum values.

### Il11 expression in smooth muscle cells activates non-canonical Il11 signaling pathways

Given that smooth muscle cells are expressed in the walls of most organs, including the vasculature, bronchi, gastrointestinal and abdominal organs, we sought to confirm the expression of *Il11* in *Il11*^SMC^ mice in specific organs. We performed western blotting across tissues harvested at 14 days after tamoxifen administration. This confirmed that Il11 protein was significantly upregulated at the protein level across all tissues tested (P_colon_ = 0.034; P_heart_ = 0.002; P_lung_ = 0.039; P_liver_ < 0.001; P_kidney_ = 0.004; and P_skin_ = 0.004; **Fig. 3** and **Supplementary Figure 3**).

**Fig 3.**
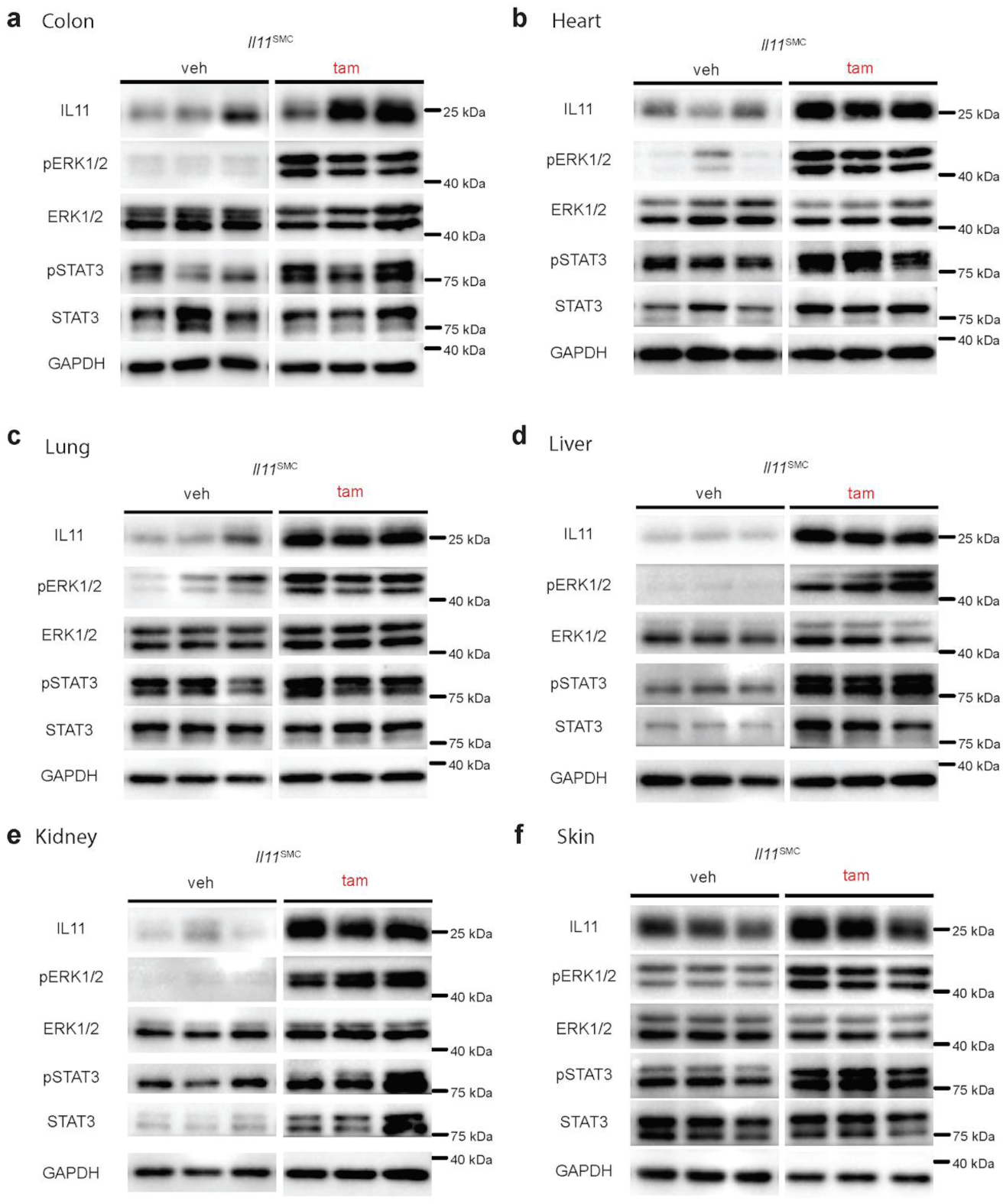
*Il11*^SMC^ mice exhibit activated ERK1/2 signaling across organs. **a-f** Immunoblots of Il11 expression, phospho-(p) and total ERK1/2 and STAT3 protein in colon, heart, liver, lung, kidney, skin tissue of *Il11*^SMC^ mice treated with vehicle or tamoxifen (*n* = 3 per group). Total and phosphorylated protein levels were quantified and normalised as detailed in **Supplementary Figure 3**. All comparisons were conducted in organs harvested from mice 14 days post-veh or tam treatment.

IL11 is a member of the IL6 family of cytokines, which are considered to signal via the Janus Kinase (JAK)/Signal Transducer and Activator of Transcription (STAT) pathway [16]. However, we recently showed that the IL11 effect, both *in vitro* in fibroblasts and *in vivo* at the tissue level, is also dependent on non-canonical signaling via extracellular signal-regulated kinase (ERK) [8–10]. To investigate both canonical and non-canonical signaling pathways after *Il11* expression, we performed western blotting of phosphorylated (p) STAT3 or ERK1/2 and total protein levels and derived indices of kinase activation by normalising phosphorylation amounts to total protein levels (**Fig 3 and Supplementary Figure 3**). At baseline, ERK was phosphorylated at low levels in most tissues except for the skin. Upon IL11 expression, we detected a strong and significant activation of ERK in all tissues (P_colon_ = 0.002; P_heart_ = 0.004; P_lung_ = 0.049; P_liver_ = 0.056; P_kidney_ = 0.001; and P_skin_ < 0.001; **Supplementary Figure 3)**. STAT3 phosphorylation was unchanged in the heart, lung and liver but was elevated in the colon and skin (P = 0.05 and 0.001 respectively; **Supplementary Figure 3**). In contrast, total levels of STAT3 appeared to be increased in the liver and kidney of tam-treated *Il11*^SMC^ animals (**Figure 3d and e**). Overall, while both pathways were affected, ERK signaling was consistently activated across tissues tested whereas STAT3 was not.

### IL11 destroys tissue integrity and promotes collagen deposition

To investigate the effect of *Il11* expression in SMCs on tissue composition beyond the colon, we performed histological analyses of the heart, lung, liver, kidney and skin. Masson’s trichrome staining was used to visualize collagen and quantify extracellular matrix deposition. In the heart, we observed collagen deposition in the perivascular region (P = 0.002; **Fig 4a, b**). We also observed vascular hypertrophy (P = 0.019; **Fig 4c**) and mild ventricular hypertrophy in the absence of dilatation (data not shown). Hydroxyproline assay of the whole heart confirmed cardiac fibrosis (P = 0.026; **Fig 4d**). In the lung, Ashcroft scores of pulmonary histological images showed lung damage after tam-induced *Il11* expression (P < 0.001; **Fig 4e, f**). Masson’s trichrome staining indicated elevated collagen expression throughout the lung in *Il11*^SMC^ mice and pulmonary fibrosis was confirmed by the hydroxyproline assay (P = 0.001; **Fig 4g**).

**Fig 4.**
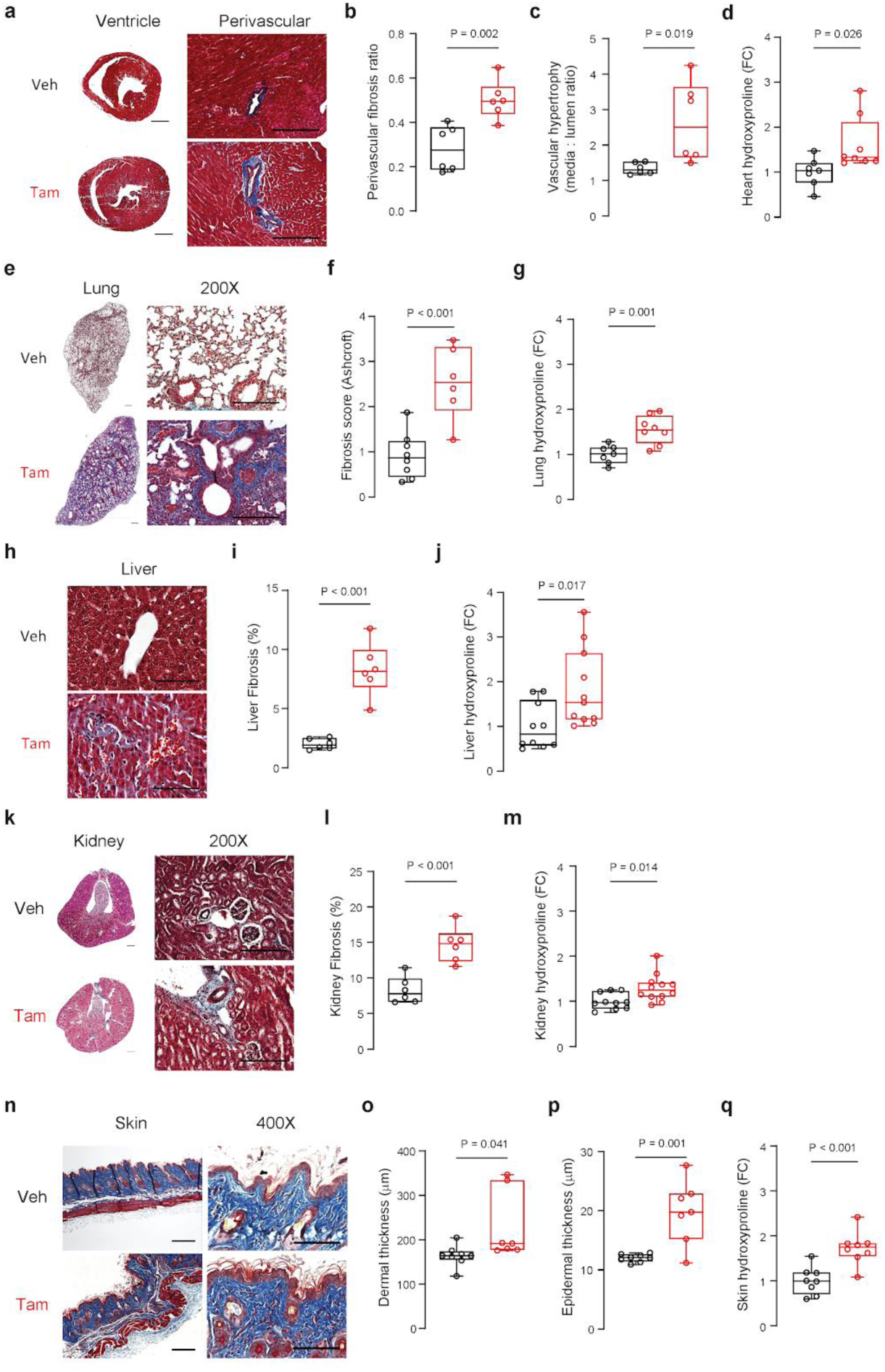
*Il11* expression in smooth muscle cells causes fibrosis across organs. **a** Representative Masson’s trichrome stained mid-ventricle sections of the heart harvested at 14 days post-veh or tam initiation (left) and 200X magnification images demonstrating perivascular fibrosis (right). Scale bars for mid-ventricle sections and 200X magnification denote 500 µm and 200 µm respectively. **b** Perivascular fibrosis quantification of histological images (200X) from veh- and tam-treated *Il11*^SMC^ mice (*n* = 6 per group). **c** Vascular hypertrophy quantification of veh- and tam-treated *Il11*^SMC^ mice (*n* = 6 per group). **d** Total collagen content in the heart assessed by hydroxyproline assay and shown as fold change (FC) of veh- and tam-treated *Il11*^SMC^ mice (*n* = 7-8 per group). **e** Representative Masson’s trichrome stained whole lung sections (left) and 200X magnification images (right). Scale bars for whole lung sections and 200X magnification denote 500 µm and 200 µm respectively. **f** Pulmonary fibrosis quantification as assessed by the Ashcroft score (*n* = 6-8 per group). **g** Total collagen content in the lung assessed by hydroxyproline assay as above (*n* = 7-8 per group). **h** Representative Masson’s trichrome stained liver sections taken at 400X magnification demonstrating perisinusoidal fibrosis. Scale bar at 400X magnification indicates 100 µm. **i** Fibrosis quantification of liver sections (400X magnification) from veh- and tam-treated *Il11*^SMC^ mice (*n* = 6 per group). **j** Total collagen content in the liver assessed by hydroxyproline assay as above (*n* = 10-11 per group). **k** Representative Masson’s trichrome stained cross-section of the kidney (left) and 200X magnification images (right). Scale bars for the cross-section of the kidney and 200X magnification denote 500 µm and 200 µm respectively. **l** Fibrosis quantification of kidney sections (200X magnification) from veh- and tam-treated *Il11*^SMC^ mice (*n* = 6 per group). **m** Total collagen content in the kidney assessed by hydroxyproline assay as above (*n* = 10-11 per group). **n** Representative Masson’s trichrome stained section of the dorsal skin at 100X magnification (left) and at 400X magnification (right). Scale bar at 100X and 400X magnification represents 200 µm and 100 µm respectively. **o** Dermal and **p** epidermal thickness of the dorsal skin. **q** Total collagen content in the skin assessed by hydroxyproline assay as above (*n* = 10-11 per group). All comparisons were conducted in organs harvested from mice 14 days post-veh and tam treatment. Statistical analyses by two-tailed unpaired t-test; data shown as median ± IQR, whiskers representing the minimum and maximum values.

The effect of *Il11* expression on the liver was overall mild and characterized by perisinusoidal fibrosis (**Fig 4h-j**). Renal tissue structure was also affected only mildly, with limited fibrosis occurring around the blood vessels (**Fig 4k-m**). The effect of IL11 on the skin of tam-treated *Il11*^SMC^ animals was more profound and both the dermal and epidermal thickness was significantly increased (**Fig 4n-p**; P = 0.041 and P = 0.001 respectively). Dorsal skin sections showed that epidermal and dermal cell infiltrates were increased and the adipose tissue layer in the hypodermis was largely depleted. Confirming Masson’s trichrome staining of skin sections, we observed increased collagen deposition in the skin of tam-treated *Il11*^SMC^ using the hydroxyproline assay (P < 0.001; **Fig 4q**).

### IL11 secretion from smooth muscle cells drives fibrogenic gene expression

To extend the observation of multi-organ fibrosis from our histology studies, we assessed the RNA expression of fibrogenic genes. Reverse transcription-polymerase chain reaction (RT-PCR) was performed using RNA from colonic, ventricular, pulmonary, hepatic, renal and skin tissue of vehicle- or tam-treated *Il11*^SMC^ mice. Collagen, type I, alpha 1 (*Col1a1*) RNA was significantly upregulated in all tissues (P_colon_ = 0.005; P_heart_ = 0.005; P_lung_ < 0.001; P_liver_ = 0.016; P_kidney_ = 0.022; P_skin_ = 0.016; **Fig 5**), confirming the effect of *Il11* expression on global organ fibrosis that we observed on the protein level (**Fig 4**). Additional markers for fibrosis such as collagen, type I, alpha 2 (*Cola1a2*), collagen, type III, alpha 1 (*Col3a1*), fibronectin 1 (*Fn1*), tissue inhibitor of metalloproteinase 1 (*Timp1*) and matrix metallopeptidase 2 (*Mmp2*) were also assessed via RT-PCR (**Fig 5a-f**). These genes were elevated in most tissues of tam-treated *Il11*^SMC^ mice. *Timp1* transcripts were significantly upregulated in the heart (P < 0.001), lung (P = 0.004), liver (P = 0.003), kidney (P = 0.003) and skin (P = 0.017), which is a recognised feature of pathological ECM remodeling [17].

**Fig 5.**
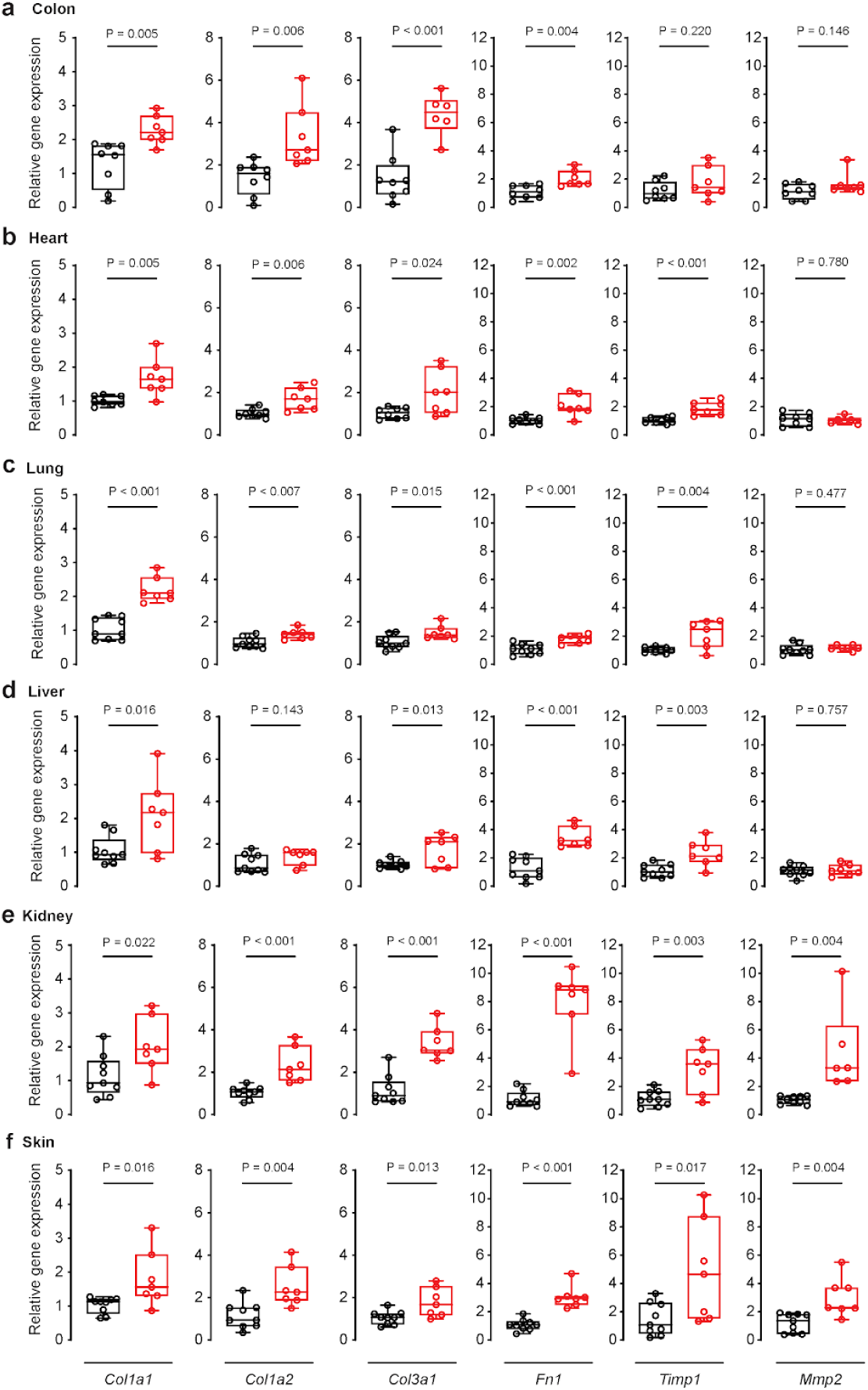
Relative gene expression of fibrogenic genes in organs from tam-treated *Il11*^SMC^ mice. Relative mRNA expression of collagen type 1a1 (*Col1a1*), type 1a2 (*Col1a2*), type 3a1 (*Col3a1*), fibronectin-1 (*Fn1*), tissue inhibitor of metalloproteinase 1 (*Timp1*) and matrix metalloproteinase 2 (*Mmp2*) normalized to Glyceraldehyde 3-phosphate dehydrogenase (*Gapdh*) expression in the **a** colon, **b** heart, **c** lung, **d** liver, **e** kidney and **f** skin. All comparisons were conducted 14 days post-veh (black) and tam (red) initiated mice. Statistical analyses by two-tailed unpaired t-test; data expressed as median ± IQR, whiskers representing the minimum and maximum values.

### IL11 secreted from smooth muscle cells causes widespread inflammation

In addition to fibrosis, SMC-driven diseases are often characterized by tissue inflammation. To better understand whether IL11 secretion from SMCs can contribute to this pathology, we performed RT-PCR experiments of inflammatory marker genes across multiple tissues. Interleukin 6 (IL6) also signals via gp130, similar to IL11, but its specific IL6 receptor subunit is expressed on a different subset of cells, most of which belong to the immune system [8]. IL6 is also a well-established therapeutic target for inflammatory diseases such as rheumatoid arthritis [18]. Upon tam-induced *Il11* expression in *Il11*^SMC^ mice, we found *Il6* mRNA to be significantly upregulated across all tissues tested (P_colon_ = 0.001; P_heart_ < 0.001; P_lung_ = 0.015; P_liver_ = 0.007; P_kidney_ < 0.001; and P_skin_ = 0.003; **Fig 6)**.

**Fig 6.**
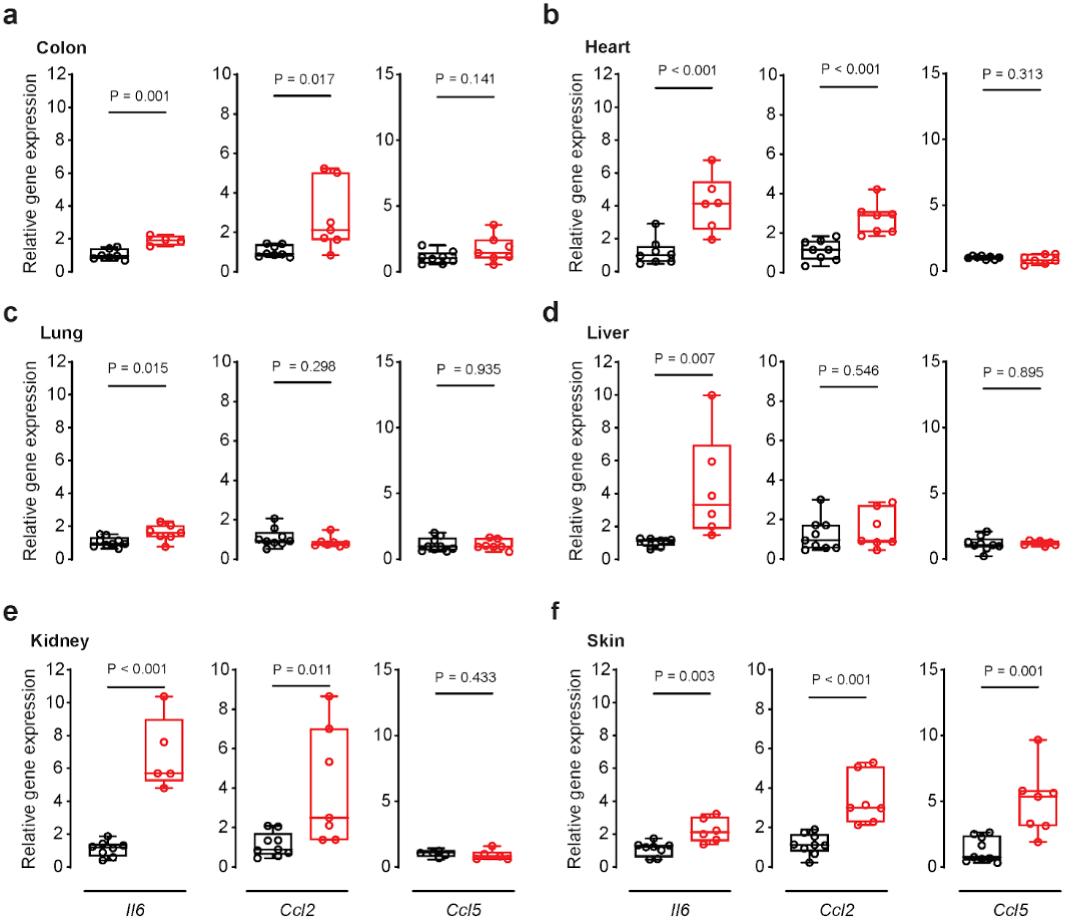
Relative gene expression of inflammatory genes in organs from tam-treated *Il11*^SMC^ mice. **a-f** Relative mRNA expression of interleukin 6 (*Il6*), C-C motif chemokine ligand 2 (*Ccl2*), C-C motif chemokine ligand 5 (*Ccl5*) normalized to Glyceraldehyde 3-phosphate dehydrogenase (*Gapdh*) expression in the colon, heart, lung, liver, kidney and skin respectively. All comparisons were conducted in 14 days post-veh (black) and tam (red) initiated mice. Statistical analyses by two-tailed unpaired *t*-test; data expressed as median ± IQR, whiskers representing the minimum and maximum values.

In the colon, we also detected increased RNA expression of the inflammatory chemokine C-C motif chemokine ligand 2 (*Ccl2*) (P = 0.017), whereas C-C motif chemokine ligand 5 (*Ccl5*) was not significantly elevated but trended upwards (P = 0.141). Interestingly, these inflammatory chemokines are upregulated in the colonic mucosa of IBD patients [19,20]. However, *CCL2* transcripts, and not *CCL5* transcripts, were found to be expressed in vessel-associated cells such as SMCs in IBD [19]. Given that *Il11*^SMC^ mice express *Il11* in SMCs, it is consistent that the transcript expression of the chemokine expressed in this particular cellular niche in the colon is most affected. In the skin, all three inflammatory markers tested were highly upregulated. This points to an inflammatory gene expression signature in the skin that is reminiscent of that seen in systemic sclerosis, since *IL6, CCL2*, and *CCL5* are elevated in the serum of patients [21,22]. Of note, *CCL2* levels were correlated with the extent of skin fibrosis in systemic sclerosis, a pathogenic feature also triggered by IL11 expression in SMCs (**Fig. 4**, **5**) [21].

### Fibroblast-selective expression of Il11 recapitulates the colonic inflammatory phenotype seen in Il11^SMC^ mice

We have previously described a model of *Il11* expression in fibroblasts (*Il11*^Fib^) that drives fibrosis in the heart, kidney, and lung [8,9]. To examine further the effect of *Il11* expression in stromal cells on the colon, we studied colonic phenotypes in this second model of *Il11* expression from the stromal niche (**Fig 7a**). Gross examination of the gastrointestinal tract of *Il11*^Fib^ mice revealed macroscopic appearances consistent with inflammation of the colon to a similar extent as in *Il11*^SMC^ mice (data not shown). The total gastrointestinal gut length of *Il11*^Fib^ mice was unchanged overall but the colon length alone was reduced (P = 0.030; **Fig 7b-c**), which is a feature of experimental colitis in mice [23]. Inflammation of the gut was apparent in the *Il11*^Fib^ model as fecal calprotectin was significantly elevated (P = 0.003). In this model, as compared to *Il11*^SMC^ mice, we detected *Il6* but not *Ccl2* or *Ccl5*, upregulation in the colon (**Fig 7d, e**). Histological examination revealed marked colonic dilation and increased SMC thickness (**Fig 7f, g**). In contrast to the *Il11*^SMC^ model of *Il11* expression, colonic fibrosis as determined by histology, hydroxyproline assay or ECM gene expression, was not significantly different between tam-treated *Il11*^Fib^ and controls (data not shown). Taken together, fibroblast-driven *Il11* expression recapitulates primarily the SMC-driven inflammatory, but not the fibrotic, phenotype in the mouse.

**Fig 7.**
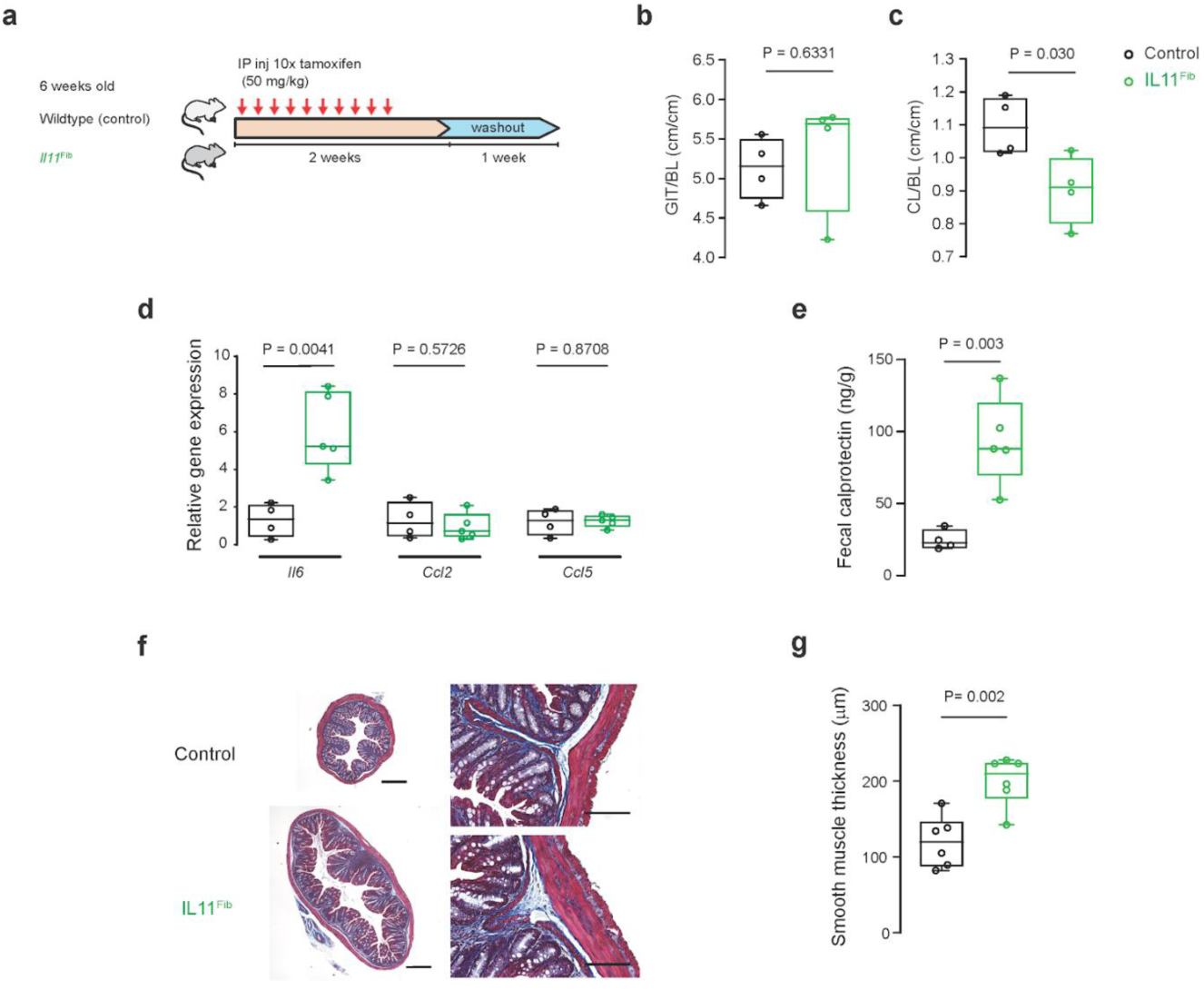
Mice with fibroblast-specific *Il11* expression develop inflammatory bowel disease. **a** Schematic diagram demonstrating tamoxifen (tam) injection procedure in 6-week-old *Il11*^Fib^ and wildtype (control) littermates. **b-c** Indexed GIT length in reference to body length (BL) was unchanged in *Il11*^Fib^ mice but indexed colon length was markedly reduced as compared to controls (*n* = 4 per group). **d** Expression of inflammatory genes (*Il6, Ccl2* and *Ccl5*) in the colon tissue of *Il11*^Fib^ mice as compared to controls (*n* = 4-5 per group). **e** Fecal calprotectin in stool samples collected from *Il11*^Fib^ and control mice (*n* = 4-5 per group) as assessed by ELISA. **f** Representative cross-sections of the colon of *Il11*^Fib^ and control mice stained with Masson’s Trichrome (left) and at 200X magnification (right) (*n* = 6 biological replicates). Scale bars indicate 500 µm and 200 µm respectively. **g** Thickness of the smooth muscle layer (muscularis propria) in tam-treated *Il11*^Fib^ mice compared to controls (*n* = 6 per group). All comparisons were conducted in 21 days post-tam initiation in control (black) and *Il11*^Fib^ (green) mice. Statistical analyses by two-tailed unpaired *t*-test; data expressed as median ± IQR, whiskers representing the minimum and maximum values.

## Discussion

In humans, *IL11* is highly upregulated in the colonic mucosa of patients with either ulcerative colitis or Crohn’s disease who do not respond to anti-TNF therapy, with recent single cell RNA-seq studies localizing IL11 to inflammatory mucosal stromal cells [24–26]. To better understand the effect of IL11 in the colon, recombinant human IL11 has been used in rodent models of IBD [27–30] and it was suggested that IL11 may have a protective role in the bowel. However, a caveat with these studies is that human IL11 was administered to rodents despite the fact that human IL11 does not activate some types of mouse cells [8]. Hence, there is a need to assess the effects of species-specific IL11 in the mouse, which we undertook in this study by expressing murine *Il11* in SMCs or fibroblasts in adult mice.

To enable our studies, we developed the *Il11*^SMC^ mouse as a tool to study the effect of murine IL11 secreted from SMCs, an established source of IL11 in the vasculature, airway, and colon [11,31,32]. Surprisingly, expression of *Il11* in SMCs was sufficient to induce severe colonic inflammation and rectal prolapse within 3 days, which was followed by early mortality in *Il11*^SMC^ animals. We also documented increased colonic muscle thickness, which is a characteristic of the dextran sulphate sodium-induced colitis model [33].

We explored further the IL11 effect in the bowel using an additional model that expresses mouse *Il11* in a second stromal cell type: the fibroblast. This complementary model also develops severe diarrhea and inflammation of the colon, reinforcing the data generated in the *Il11*^SMC^ mice. These data show that *Il11* expression alone in stromal cells is sufficient to cause an IBD phenotype and challenges the earlier data, based on the use of human IL11 in the mouse. The effect of IL11 on the vasculature will be discussed elsewhere. Considered together with patient studies that show IL11 to be highly upregulated in the colonic mucosa of patients with ulcerative colitis or Crohn’s disease [24–26], our results highlight IL11 as a promising therapeutic target for IBD, particularly in the context of anti-TNF therapy resistance.

## Materials and Methods

### Mouse models

All experimental procedures were approved and conducted in accordance to the SingHealth Institutional Animal Care and Use Committee (IACUC). All mice were from a C57BL/6JN genetic background and they were bred and housed in the same room and provided food and water *ad libitum*.

#### Smooth muscle-specific Il11 transgenic model

To direct transgene expression in smooth muscle cells, we crossed the heterozygous *Rosa26-Il11* (Gt(ROSA)26Sor^tm1(CAG-Il11)Cook^) mouse [8] to the hemizygous *SMMHC-CreERT2* (B6.FVB-Tg(Myh11-cre/ERT2)1Soff/J) mouse [15] available from the Jackson Laboratory (031928 and 019079 respectively) to generate double heterozygous *SMMHC-CreERT2:Rosa26-IL11* offspring (referred to here as *Il11*^SMC^ mice). Only male *Il11*^SMC^ mice were utilized as the *Myh11-Cre/ERT2* transgene is inserted on the Y chromosome. To induce Cre-mediated *Il11* transgene induction, six week old *Il11*^SMC^ mice were intraperitoneal injected with 3 doses of 50 mg kg^-1^ tamoxifen (tam; T5648, Sigma Aldrich) or an equivalent volume of corn oil vehicle (veh; C8267, Sigma Aldrich) for a week. Single hemizygous *SMMHC-CreERT2* littermates were designated as controls (referred to as *Cre*^SMC^). Mice were euthanized at 14 days following the first injection.

For genotyping of mice genomic DNA, we performed polymerase chain reaction (PCR) on the tail biopsies which were obtained at the time of weaning. Genotyping was conducted in two sequential PCRs, for *Myh11-Cre* and *Rosa26-Il11* genes separately. Agarose gel electrophoresis was subsequently conducted to confirm the respective product sizes for genotyping. Genotyping primer sequences are listed in **Supplementary Table 1**.

#### Fibroblast-specific Il11 transgenic model

To model fibroblasts-selective secretion of IL11 *in vivo*, we crossed the heterozygous *Rosa26-IL11* mice with *Col1a2-CreER* mice [34] to generate double heterozygous *Col1a2-CreER:Rosa26-Il11* mice (referred to as *Il11*^Fib^) [9]. For Cre-mediated *Il11* transgene induction, *Il11*^Fib^ mice were intraperitoneal injected with 50 mg kg^-1^ tamoxifen at 6 weeks of age for 10 consecutive days and the animals were sacrificed on day 21. Wildtype littermates (designated as control) were injected with an equivalent dose of tamoxifen for 10 consecutive days as controls. Both female and male mice were used.

Colon length was measured from the caecum to the anus. The most distal half was taken for histology and the adjacent part was portioned and immediately snap frozen in liquid nitrogen for downstream molecular work (hydroxyproline assay, western blot analysis and quantitative polymerase chain reaction assessment). The excised heart was halved from the base to mid ventricle for histology and the remainder separated into 3 portions for molecular work. The left lung was isolated for histology and the right lung separated into 3 portions for molecular work. The right lobe of the liver was excised for histology and the left lobe separated into 3 portions for molecular work. The left kidney was fixed for histology and the right kidney separated in thirds for molecular work. The dorsal skin was harvested and halved for histology and molecular work.

### Hydroxyproline assay

The amount of total tissue collagen was quantified using colorimetric detection of hydroxyproline using the Quickzyme Total Collagen assay kit (Quickzyme Biosciences) performed according to the manufacturer’s protocol. All samples were run in duplicate and absorbance at 570 nm was detected on a SpectraMax M3 fluorescence microplate reader using SoftMax Pro version 6.2.1 software (Molecular Devices).

### Fecal calprotectin (S100A8/A9) levels

To characterize inflammation in the gut, we investigated levels of fecal calprotectin in the *Il11*^SMC^ and *Il11*^Fib^ mice using the mouse S100A8/A9 heterodimer duoset ELISA kit (DY8596-05, R&D systems). Calprotectin is a biomarker for inflammatory activity and has been clinically applied as a diagnostic tool for inflammatory bowel diseases [35,36]. Stool samples were collected in a 1.5 ml tube and diluted with 50x (weight per volume) of extraction buffer (0.1 M Tris, 0.15 M NaCl, 1.0 M urea, 10 mM CaCl2, 0.1 M citric acid monohydrate, 5 g/l BSA (pH 8.0)) with the assumption of fecal density to be 1 g/ml. Samples were homogenized until no large particles were present. Homogenate was transferred into a fresh tube and further centrifuged at 10,000 g at 4 °C for 20 minutes. The supernatant was assessed for S100A8/A9 levels by ELISA as per the manufacturer’s instructions.

### RT-qPCR

Total RNA was extracted from snap-frozen tissues using RNAzol RT (R4533, Sigma-Aldrich) followed by Purelink RNA mini kit (12183025, Invitrogen) purification. The cDNA was prepared using iScript cDNA synthesis kit (1708891, Bio-Rad) as per the manufacturer’s instructions. Quantitative RT-PCR gene expression analysis was performed on duplicate samples using fast SYBR green (Qiagen) technology using the ViiA 7 Real-Time PCR System (Applied Biosystem). RT-qPCR primers are listed in **Supplementary Table 2**. Expression data were normalized to *Gapdh* mRNA expression levels and the 2^-ΔΔCT^ method was used to calculate the fold change.

### Immunoblotting

Western blots were carried out on total protein extracts from mouse tissues. Frozen tissues were homogenized and lyzed in radioimmunoprecipitation assay (RIPA) buffer containing protease and phosphatase inhibitors (Roche) followed by centrifugation. Equal amounts of protein lysates were separated by SDS-PAGE, transferred onto PVDF membrane and immunoblotted for pERK1/2 (4370, CST), ERK1/2 (4695, CST), pSTAT3 (4113, CST), STAT3 (4904, CST), GAPDH (2118, CST) and IL11 (X203, Aldevron). Proteins were visualized with appropriate secondary antibodies anti-rabbit HRP (7074, CST) and anti-mouse HRP (7076, CST).

### Histology

Tissues from *Il11*^SMC^ and *Il11*^Fib^ mice were fixed in 10% neutral-buffered formalin for 24-48 hours, tissue processed and paraffin-embedded. Sections were obtained at 5 µm and stained with Masson’s trichrome staining for collagen. Brightfield photomicrographs of the sections were randomly captured by a researcher blinded to the treatment groups using the Olympus BX51 microscope and Image-Pro Premier 9.2 (Media Cybernetics).

Photomicrographs of the colon taken at 200X magnification were used to calculate muscle wall thickness. The distance between the inner and outer circumference of the muscularis propria was measured using the incremental distance tool at a calibrated step size of 25 µm on Image-Pro Premier 9.2 (Media Cybernetics). A total of 75 to 250 measurements across three to five photomicrographs per section were taken and averages reported per photomicrograph. Muscle thickness was reported as an average across 3 cross-sections of the colon per animal.

Photomicrographs of the dorsal skin were captured in 3 fields per section at 100X magnification and used to calculate epidermal and dermal thickness. The epidermis was measured from the stratum basale to the stratum granulosum using hand-drawn line segments on Image-Pro Premier 9.2 (Media Cybernetics). The dermis was measured from the dermal-epidermal junction to the hypodermis. Measurements were recorded using the incremental distance tool at a calibrated step size of 50 µm on Image-Pro Premier 9.2 (Media Cybernetics). A total of 75 to 200 measurements across three photomicrographs per section were taken and averages reported per photomicrograph. Overall epidermal and dermal thickness was reported as an average across the 3 fields per animal.

Fibrosis quantification was conducted as referenced [37]. Color deconvolution version 1.5 plugin using the Masson Trichrome vector on ImageJ (version 1.52a, NIH) and thresholding was applied for area quantification. Perivascular fibrosis was measured as a ratio of the fibrosis area to the vessel area. Vascular hypertrophy was quantified as the ratio of media wall area to the lumen area.

### Statistical analysis

Data are presented as mean ± standard deviation or median ± range as stated in figure legends. Statistical analyses were performed on GraphPad Prism 8 software (version 8.1.2). Outliers (ROUT 2%, GraphPad Prism software) were removed prior to analyses. Comparison of survival curves was analyzed with the log-rank Mantel-Cox test. Bodyweight progression was analyzed with two-way ANOVA with Sidak multiple comparisons. A comparison of mice strains for all other parameters was analyzed with a two-tailed unpaired *t*-test. The criterion for statistical significance was established at P < 0.05.

## Acknowledgements

The authors would like to acknowledge B.L. George, E. Khin, M. Wang, J. Tan for their technical expertise and support.

## Author contributions

Conceived and designed the experiments: WWL BN SAC SS. Performed the experiments WWL BN AW CX LS NK SYL XK SL. Analyzed the data: WWL BN. Manuscript writing, review and editing: WWL BN SAC SS.

## Supplementary Tables

**Supplementary Table 1.**
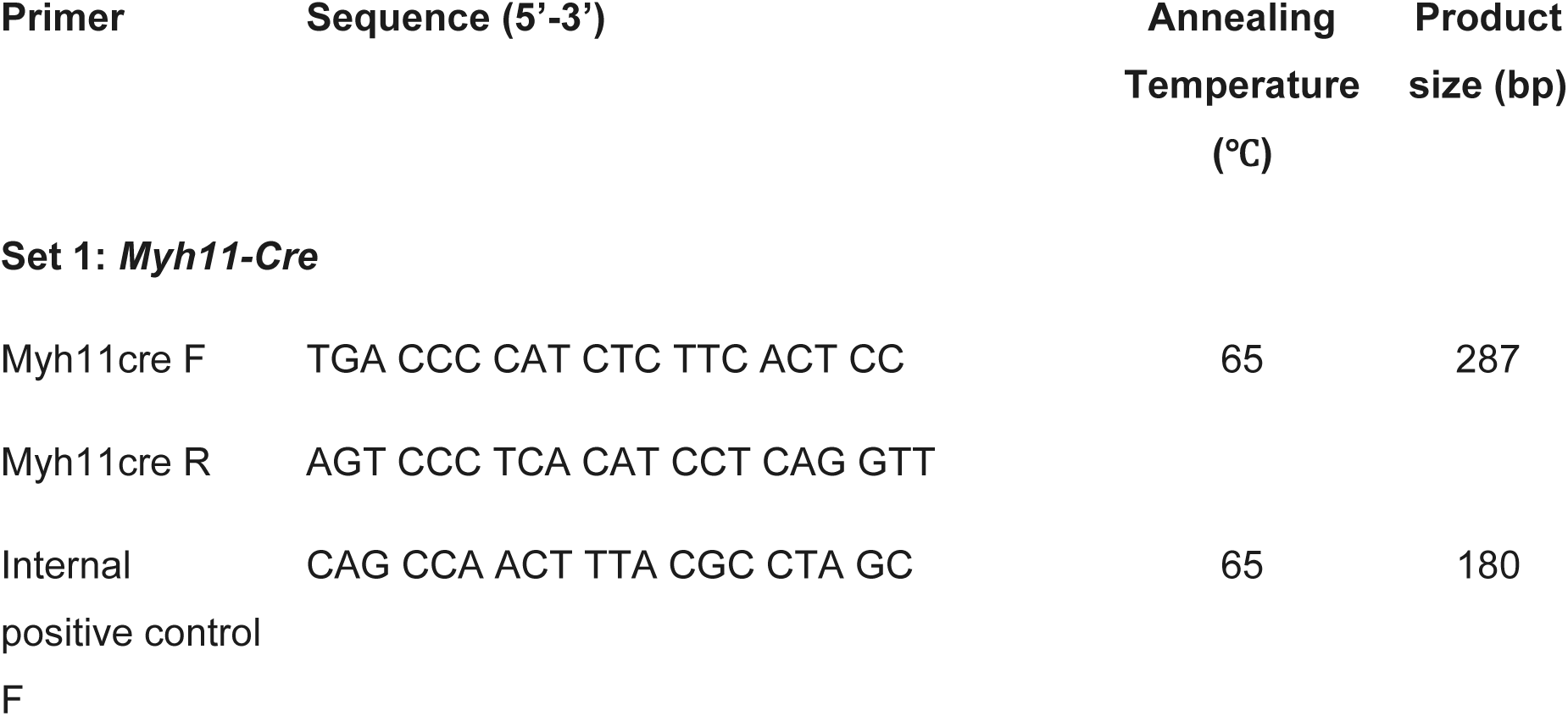
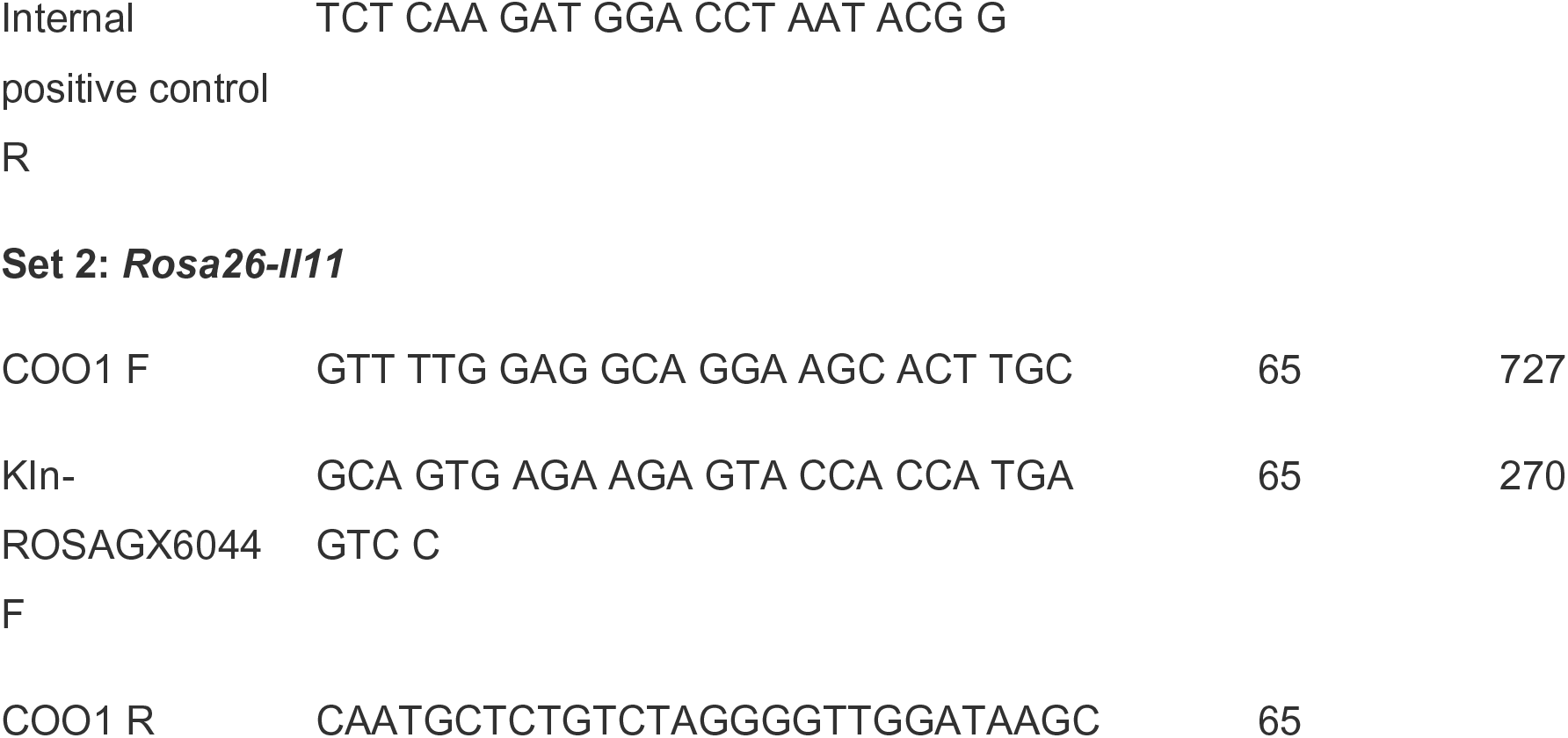
Genotyping primers.

**Supplementary Table 2.**
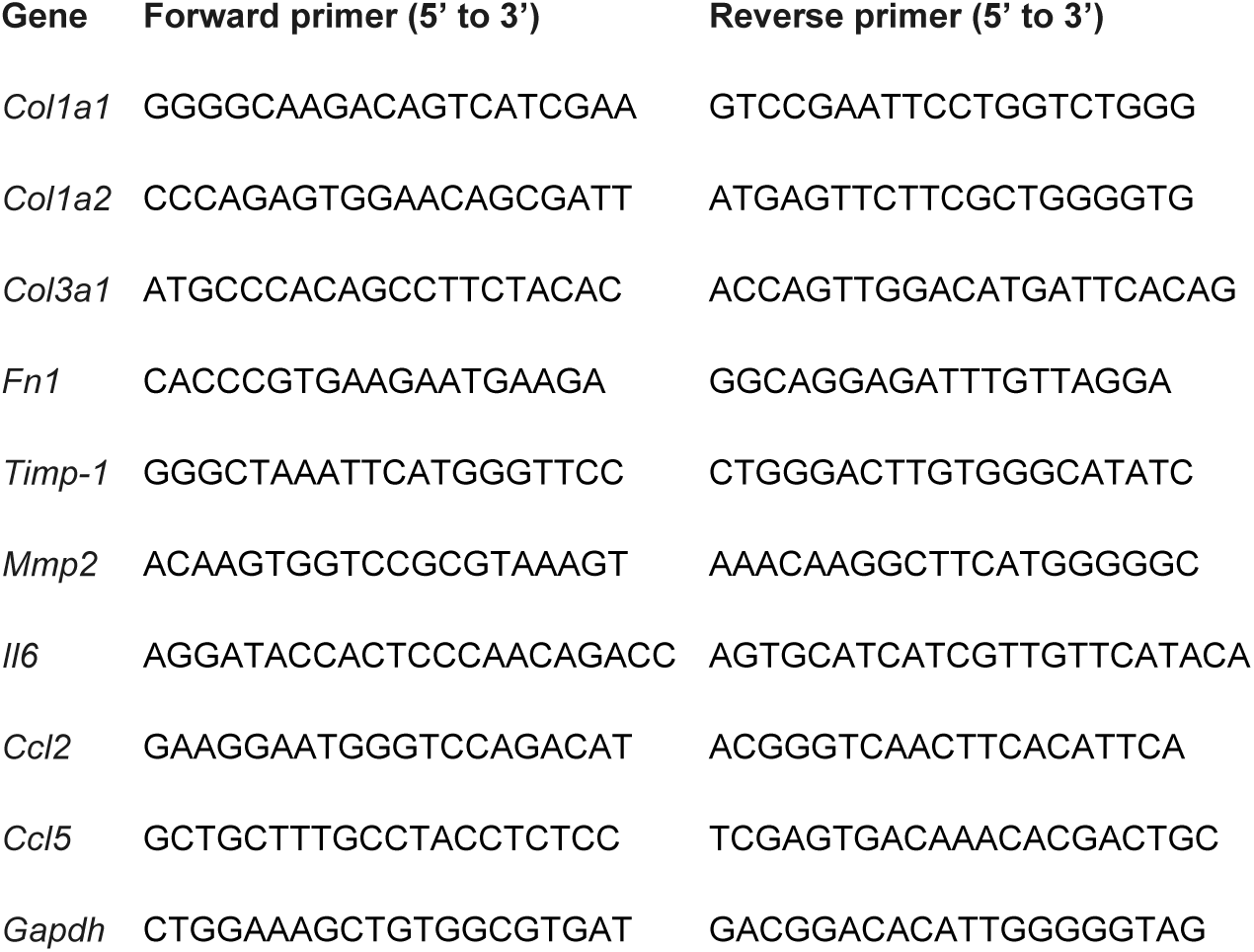
RT-qPCR primers.

## Supplementary Figures

**Supplementary Figure 1.**
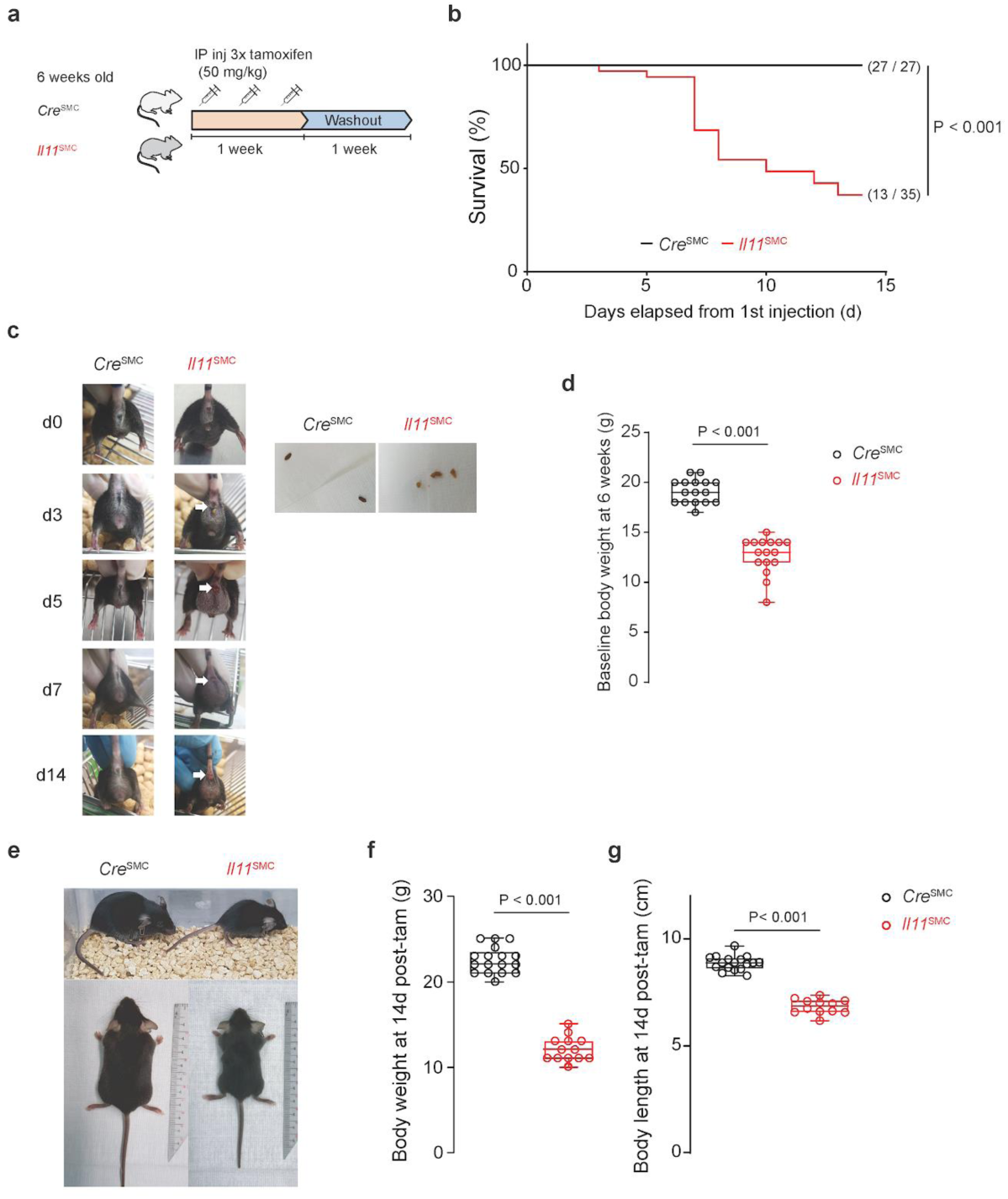
Comparison of tamoxifen-treated *Il11*^SMC^ and *Cre*^SMC^ mice. **a** Schematic diagram demonstrating the tamoxifen (tam) injection procedure in 6-week-old *Il11*^SMC^ and *Cre*^SMC^ mice. **b** Survival curve of tam-treated *Il11*^SMC^ (*n* = 35) compared to *Cre*^SMC^ mice (*n* = 27) mice from 1st injection starting at 6 weeks of age. Survival curves were compared with the log-rank Mantel-Cox test. **c** Representative images of the *Cre*^SMC^ and *Il11*^SMC^ mice before (d0) and up to 14 days (d14) post-tam initiation (left). Note the presence of pale and loose stools in *Il11*^SMC^ mice (right). The presence of rectal prolapse is indicated with white arrows. Tam-treated *Il11*^SMC^ images presented here are different from Fig. 1c. Images were not taken to scale. **d** Baseline body weight of 6-week-old *Cre*^SMC^ and *Il11*^SMC^ mice before induction (*n* = 16 per group). Statistical analyses by two-tailed unpaired t-test; data expressed as median ± IQR, whiskers representing the minimum and maximum values. **e** Representative images of *Cre*^SMC^ and *Il11*^SMC^ mice at d14 post-Tam initiation. **f-g** Collated body weights and body lengths of tam-treated *Cre*^SMC^ and *Il11*^SMC^ mice measured at d14 post-Tam initiation. (*n* = 12-17 per group). Statistical analyses by two-tailed unpaired t-test; data expressed as median ± IQR, whiskers representing the minimum and maximum values.

**Supplementary Figure 2.**
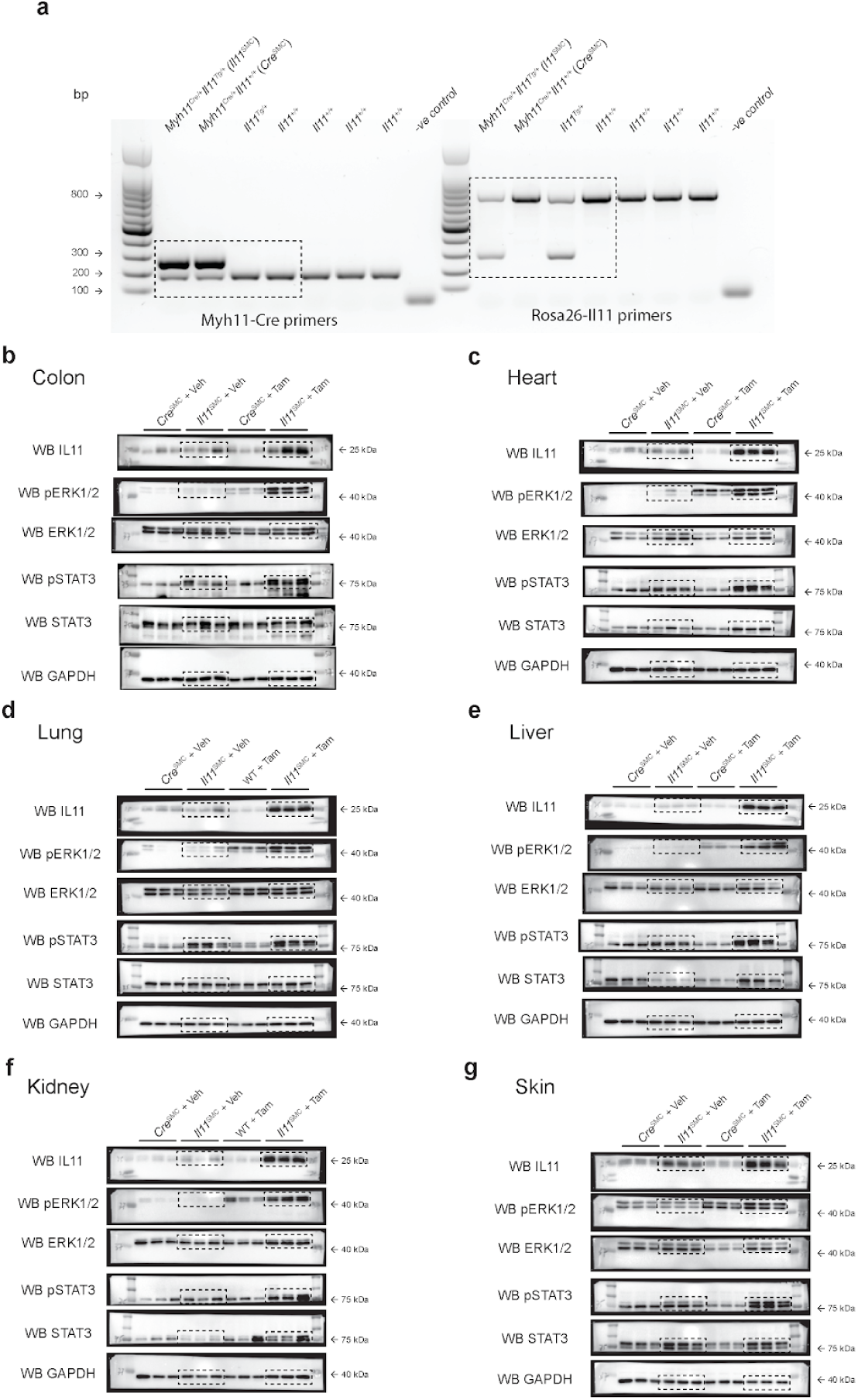
Uncropped blots for PCR genotyping and immunoblots of different organs from *Cre*^SMC^ and *Il11*^SMC^ mice treated with either tamoxifen (Tam) or vehicle (Veh) for IL11 and downstream ERK1/2 and STAT3 activation. **a** PCR products of DNA extracted from tail biopsies of 21-day-old mice by use of set primers for *Myh11-Cre* (left) and *Rosa26-Il11* (right) (primers as listed in Table S1) and analyzed by agarose gel electrophoresis. Dashed boxes indicate cropped blots used in Fig 1c. **b-g** Immunoblots of the colon, heart, lung, liver, kidney and skin tissues in *Cre*^SMC^ and *Il11*^SMC^ mice treated with Tam or Veh for IL11 and ERK1/2 and STAT3 activation. Dashed boxes indicate cropped blots used in Fig 3.

**Supplementary Figure 3.**
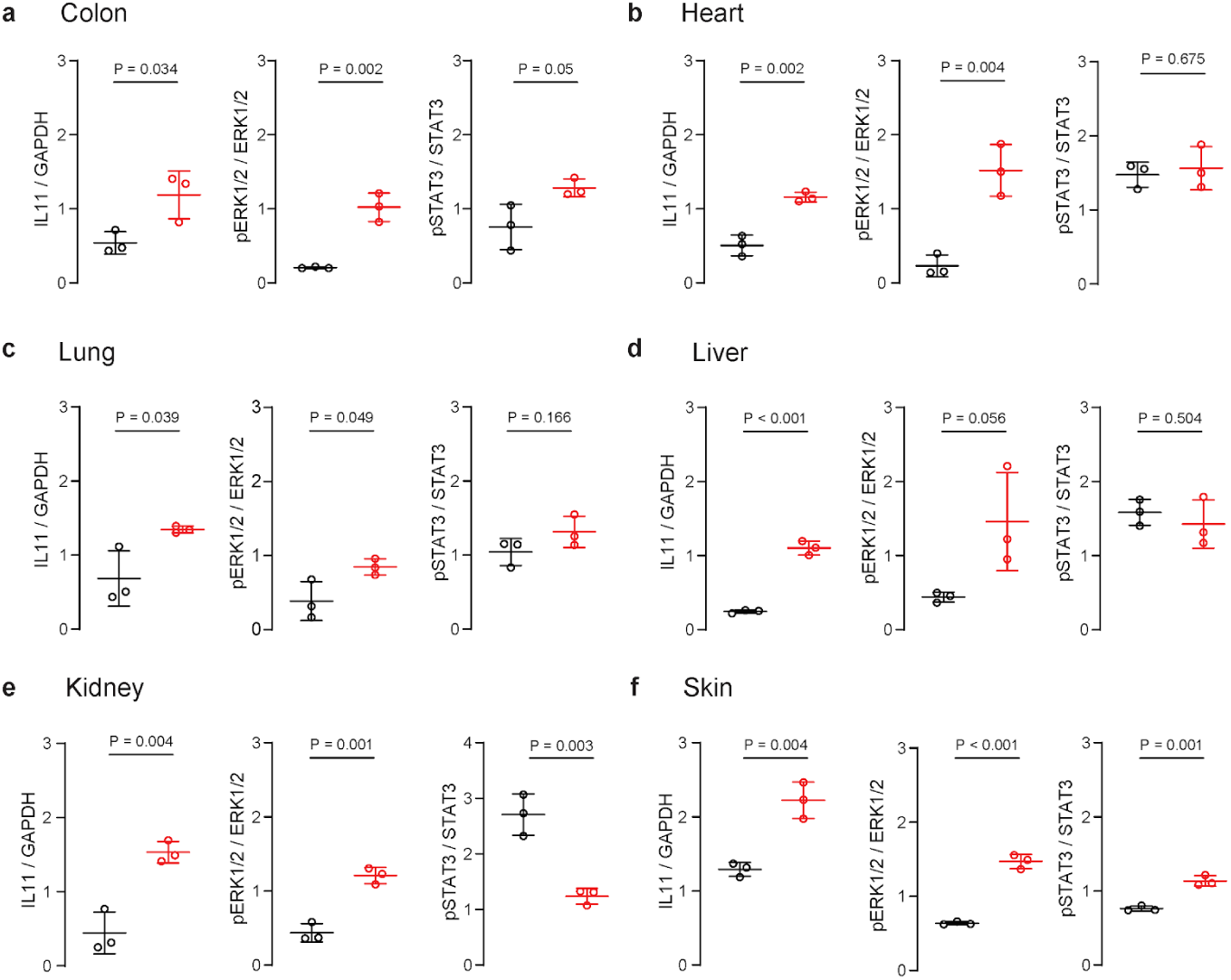
Densitometry quantification of immunoblots in Fig 2. **a-f** densitometry of immunoblots of IL11 expression, p-ERK1/2/ERK1/2 and p-STAT3/STAT3 protein in colon, heart, liver, lung, kidney, colon and skin tissue of *Il11*^SMC^ mice treated with vehicle or tamoxifen respectively. Statistical analyses by two-tailed unpaired t-test; data expressed as mean ± standard deviation.

